# Hyperactivation of extracellular signal-regulated kinase (ERK) by RAS-mediated signaling or inhibition of dual specificity phosphatase 6 (DUSP6) is associated with toxicity in lung adenocarcinoma cells with mutations in *KRAS* or *EGFR*

**DOI:** 10.1101/222505

**Authors:** Arun M. Unni, Bryant Harbourne, Min Hee Oh, Sophia Wild, William W. Lockwood, Harold Varmus

**Affiliations:** Meyer Cancer Center, Weill Cornell Medicine, New York, NY.; Department of Integrative Oncology, British Columbia Cancer Agency, Vancouver, BC.; Department of Pathology and Laboratory Medicine, University of British Columbia, Vancouver, BC.

## Abstract

We recently described the synthetic lethality that results when mutant KRAS and mutant EGFR are coexpressed in human lung adenocarcinoma (LUAD) cells, revealing the biological basis for the mutual exclusivity of *KRAS* and *EGFR* mutations in lung cancers. We have now further defined the biochemical events responsible for the toxic effects of signaling through the RAS pathway. By combining pharmacological and genetic approaches, we have developed multiple lines of evidence that signaling through extracellular signal-regulated kinases (ERK1/2) mediates the toxicity. These findings imply that tumors with mutant oncogenes that drive signaling through the RAS pathway must restrain the activity of ERK1/2 to avoid cell toxicities and enable tumor growth. In particular, a dual specificity phosphatase, DUSP6, regulates phosphorylated (P)-ERK levels in lung adenocarcinoma cells, providing negative feedback to the RAS signaling pathway. Accordingly, inhibition of DUSP6 is cytotoxic in LUAD cells driven by either mutant KRAS or mutant EGFR, phenocopying the effects of co-expression of mutant KRAS and EGFR. Together, these data suggest that targeting DUSP6 or other feedback regulators of the EGFR-KRAS-ERK pathway may offer a strategy for treating certain cancers by exceeding an upper threshold of RAS-mediated signaling.

## Introduction

Extensive characterization of cancer genomes has begun to change the classification of neoplasms and the choice of therapies^1^. The genetic profiles of most cancers are notoriously heterogeneous, often including thousands of mutations affecting genes with a wide range of credentials---from those well-known to drive oncogenic behavior to those not known to have a role in pathogenesis. Moreover, cancers continue to accumulate mutations during carcinogenesis, producing tumor subclones with selectable features such as drug resistance or enhanced growth potential^2^. Despite this heterogeneity, consistent patterns have been observed, such as the high frequency of gain-of-function or loss-of-function mutations affecting specific proto-oncogenes or tumor suppressor genes in cancers that arise in certain cell lineages. Conversely, coincident mutations in certain genes are rare, even when those genes are frequently mutated individually in specific types of cancer^3^.

Examples of these “mutually exclusive” pairs of mutations have been reported in a variety of cancers^4-8^; the mutual exclusivity has usually been attributed either to a loss of a selective advantage of a mutation in one gene after a change in the other has occurred (“functional redundancy”) or to the toxicity (including “synthetic lethality”) conferred by the coexistence of both mutations in the same cells. We recently reported that the mutual exclusivity of gain-of-function mutations of *EGFR* and *KRAS*, two proto-oncogenes often individually mutated in lung adenocarcinomas (LUADs), can be explained by such synthetic toxicity, despite the fact that products of these two genes operate in overlapping signaling pathways and might have been mutually exclusive because of functional redundancies^5^.

Support for the idea that the mutual exclusivity of *KRAS* and *EGFR* mutations is synthetically toxic in LUAD cells was based largely on experiments in which we used doxycycline (dox) to induce expression of mutant *EGFR* or *KRAS* alleles controlled by a tetracycline (tet)-responsive regulatory apparatus in LUAD cell lines containing endogenous mutations in the other gene^5^. When we forced mutual expression of the pair of mutant cells, the cells exhibited signs of RAS-induced toxicity such as macropinocytosis and cell death. In addition, we observed increased phosphorylation of several proteins known to operate in the extensive signaling network downstream of RAS, implying that excessive signaling, driven by the conjunction of hyperactive EGFR and KRAS proteins, might be responsible for the observed toxicity.

Recognizing that such synthetic toxicities might be exploited for therapeutic purposes, we have extended our studies of signaling via the EGFR-RAS axis, with the goal of better understanding the biochemical events that are responsible for the previously observed toxicity in LUAD cell lines. In the work reported here, we have used a variety of genetic and pharmacological approaches to seek evidence that identifies critical mediators of the previously observed toxicities. Based on several concordant findings, we argue that activation of extracellular signal-regulated kinases (ERK1 and ERK2), serine/threonine kinases in the EGFR-RAS-RAF-MEK-ERK pathway, is a critical event in the generation of toxicity, and we show that at least one feedback inhibitor of the pathway, the dual specificity phosphatase, DUSP6, is a potential target for therapeutic inhibitors that could mimic the synthetic toxicity that we previously reported.

## Results

### Synthetic lethality induced by co-expression of mutant KRAS and EGFR is mediated through increased ERK signaling

In previous work, we established that mutant EGFR and mutant KRAS are not tolerated in the same cell (synthetic lethality), by placing one of these two oncogenes under the control of an inducible promoter in cell lines carrying a mutant allele of the other oncogene. These experiments provided a likely explanation for the pattern of mutual exclusivity in LUAD^5^. While we documented several changes in cellular signaling upon induction of the second oncogene to produce toxicity, we did not establish if there is a node (or nodes) in the signaling network sensed by the cell as intolerable when both oncoproteins are produced. If such a node exists, we might be able to prevent toxicity by down-modulating the levels of activity; conversely, we might be able to exploit identification of that node to compromise or kill cancer cells.

To seek critical nodes in the RAS signaling pathway, we extended our previous study using the LUAD cell line we previously characterized (PC9, bearing the EGFR mutation, E746_A750del) and two additional LUAD lines, H358 and H1975. H358 cells express mutant KRAS (G12C), and H1975 cells express mutant EGFR (L858R/T790M). As in our earlier work, we introduced tet-regulated, mutant *KRAS* into these lines to regulate mutant KRAS in an inducible manner and used the same vector encoding GFP rather than KRAS as a control. This single-vector system includes rtTA expressed from a ubiquitin promoter, allowing us to induce KRAS with the addition of dox^9^.

KRAS or GFP were appropriately induced after dox treatment of these cell lines (Fig1A). To establish whether induction of a mutant *KRAS* transgene is detrimental to cells producing endogenous mutant KRAS (H358) or mutant EGFR (H1975) proteins, we cultured cell lines in dox for 7 days and measured the relative numbers of viable cells by incubating cultures with Alamar blue. As we previously showed, the number of viable PC9 cells is reduced by inducing mutant *KRAS* (Fig 1A). Similarly, when mutant KRAS was induced in either H358 or H1975 cells for seven days, we observed fewer viable cells compared to cells grown without dox or to cells in which GFP was induced (Fig 1A). These results indicate that increased activity of the RAS pathway, either in LUAD cells with an endogenous *KRAS* mutation (H358) or with an endogenous *EGFR* mutation (PC9 and H1975) is toxic to these cell lines.

**Figure 1:**
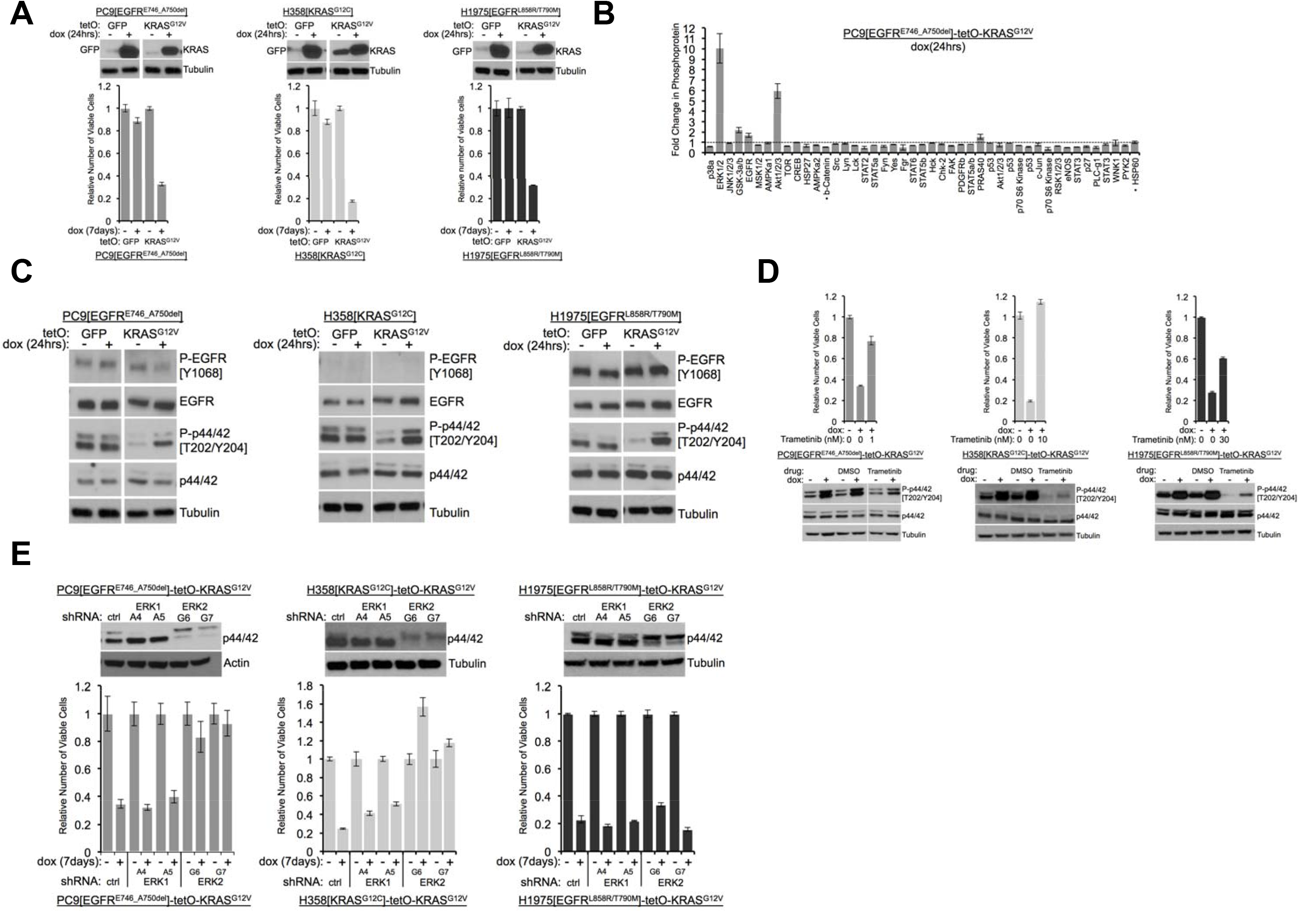
Induction of mutant KRAS reduces the numbers of viable lung cancer cells harboring KRAS or EGFR mutations, and the effects can be rescued by inhibiting ERK. (A) Reduced numbers of viable LUAD cells after activation of KRAS. Production of GFP or KRAS^G12V^ was induced by addition of 100ng/mL dox in the indicated three cell lines as described in Methods. GFP and KRAS protein levels were measured by Western blotting 24 hours later (top); the numbers of viable cells, normalized to cells grown in the absence of dox (set to 1.0), were determined by staining with Alamar blue six days later. Error bars represent standard deviations based on three replicates. (B) Induction of KRAS^G12V^ uniquely increases phosphorylation of ERK1/2 among several phosphoproteins. PC9-tetO-KRAS cells were treated with dox for 24 hrs and cell lysates incubated on an array to detect phosphorylated proteins. Fold changes of phosphorylation compared with lysates from untreated cells (set to 1.0, dotted line) to treated cells is presented from a single antibody array. Error bars are derived from duplicate spots on antibody array. The detection of HSP60 and ß-catenin are of total protein, not phosphoprotein. (C) Phosphorylation of ERK occurs early after induction of mutant KRAS. Lysates prepared as described for panel (A) were probed for the indicated proteins by western blot. Loading control is the same as in A. (D) Drug-mediated inhibition of the MEK1/2 kinases ameliorates KRAS-induced loss of viable cells. Mutant KRAS was induced with dox in the three indicated cell lines in the absence and presence of Trametinib at the indicated dose for 7 days. Cell viability was measured with Alamar blue. Error bars represent standard deviations determined from three samples grown under each set of conditions. Values are normalized to measurements of cells that received neither dox nor Trametinib (bottom). Cells were treated with dox and with or without Trametinib for 24 hours at the dose conferring rescue of numbers of viable cells. Lysates were probed for indicated proteins to confirm inhibition of MEK. (E) Reduction of ERK proteins with inhibitory RNAs protects cells from loss of viability in response to induction of mutant KRAS. LUAD cell lines, transduced with the indicated shRNA targeted against ERK1 or ERK2, were assessed for levels of ERK proteins, p42 and p44, by Western blotting (top panels). The same lines were treated with dox for 7 days and the number of viable cells measured by staining with Alamar blue. Values are normalized to numbers of viable cells of each type grown in the absence of dox (1.0), with error bars representing standard deviations among three replicates. Similar results were obtained from 2 or 3 independent experiments.

We previously documented increases in phosphorylated forms of the stress kinases, phospho-JNK (P-JNK) and phospho-p38 (P-p38), as well as in phospho-ERK (P-ERK or P-p44/42), in one of these cell lines (PC9) 72 hours after treatment with dox^5, 8^. We used a phospho-protein array to assess the status of protein activation more broadly after KRAS induction, using PC9-tetO-KRAS cells after 1 and 5 days of dox treatment (Fig 1B, Supplemental Fig 1A.). After 5 days, we again observed increases in P-JNK, P-p38, and P-ERK (Supplemental Fig 1A,), suggesting that three major branches of the MAPK pathway are activated after extended induction of mutant KRAS. In addition, several other proteins show enhanced phosphorylation at this time. At 24 hours after addition of dox, however, only P-ERK and P-AKT show a pronounced increase (Fig. 1B, right). Specifically, the stress kinases, JNK and p38, were not detected as phosphorylated proteins with the protein array. A possible interpretation of these findings is that ERK may be phosphorylated relatively soon after induction of mutant KRAS, with subsequent phosphorylation (and activation) of stress kinases and several other proteins. We also observed increased phosphorylation of ERK 24 hours after induction of mutant KRAS by western blot in all three LUAD cell lines (Fig. 1C).

These data suggest that ERK itself could be the signaling node that causes a loss of viable cells when inappropriately activated. To test this hypothesis, we used Trametinib^10^, an inhibitor of MEK, the kinase that phosphorylates ERK, to ask whether reduced levels of P-ERK levels would protect cells from the toxicity caused by induction of mutant KRAS. In all three LUAD cell lines, Trametinib completely or partially rescued the loss of viable cells caused by induction of mutant KRAS by dox (Fig 1D). We confirmed that doses of Trametinib that protected cells from the toxic effects of seven days of treatment with dox were associated with reduced levels of P-ERK after 24 hours of induction of mutant KRAS (Fig. 1D).

To extend these findings, we transduced LUAD cell lines with retroviral vectors encoding shRNAs that “knock down” expression of ERK1 or ERK2. Using two different shRNAs for each gene, as well as a non-targeted shRNA vector as control, we stably reduced the levels of ERK1 or ERK2 in the three LUAD cell lines (Fig. 1E). When PC9 and H358 lines were treated with dox to assess the effects of ERK1 or ERK2 knockdowns on the loss of viable cells, we found that depletion of ERK2, but not ERK1, rescued cells from KRAS toxicity after 7 days in dox (Fig. 1E). In H1975 cells, however, neither knockdown of ERK1 nor knockdown of ERK2 prevented KRAS-induced cell toxicity. Since Trametinib rescues the number of viable cells after induction of KRAS in H1975 cells (Fig. 1D), it seemed possible that either ERK1 or ERK2 might be sufficient to mediate RAS-induced toxicity in this line. In that case, it would be necessary to reduce the levels or the activity of both ERK proteins to rescue H1975 cells from toxicity. We tested this idea by treating dox-induced H1975-tetO-KRAS cells with SCH772984^11^, a drug that inhibits the kinase activity of both ERK1 and ERK2 (Supplementary Fig 1B). As we observed with the MEK inhibitor, Trametinib (Fig. 1D, far right), the ERK inhibitor reduces KRAS-associated toxicity in H1975 cells with concomitant reductions of P-ERK1 and P-ERK2 (Supplemental Fig 1B). Collectively, our data suggest that LUAD cell lines are sensitive to inappropriate hyperactivation of the ERK signaling node and that toxicity mediated by activation of the RAS pathway is ERK-dependent.

### DUSP6 is a major regulator of negative feedback, expressed in LUAD cells, and associated with KRAS and EGFR mutations and with high P-ERK levels

The evidence that hyperactive ERK signaling has toxic effects on LUAD cells raises the possibility that cancers driven by mutations in the RAS pathway may have a mechanism to “buffer” P-ERK levels and thereby avoid reaching a lethal signaling threshold. Genes encoding negative feedback regulators are typically activated at the transcriptional level by the EGFR-KRAS-ERK pathway to place a restraint on signaling^12^. Such feedback regulators previously implicated in the control of EGFR-KRAS-ERK signaling include the dual specificity phosphatases (DUSP1-6), the sprouty proteins (SPRY1-4) and the sprouty-related, EVH1 domain-containing proteins (SPRED1-3)^12, 13^. To begin a search for possible negative regulators of RAS-mediated signaling in LUAD cells driven by mutations in either *KRAS* or *EGFR*, we asked whether mutations in either protooncogene would up-regulate one or multiple members of these families of regulators, based on the assumption that such proteins might constrain P-ERK levels, leading to optimal growth without cytotoxic effects.

To search for potential negative regulators specifically involved in LUAD, we compared concentrations of RNAs from *DUSP, SPRY* and *SPRED* gene families in tumors with and without mutations in either *KRAS* or *EGFR*, using RNA-seq data from The Cancer Genome Atlas (TCGA)^14^ (Fig 2A). *DUSP6* was the only negative feedback regulatory gene with significantly different levels of expression when we compared tumors with mutations in either *KRAS* or *EGFR* with tumors without such mutations (Bonferoni corrected p<0.01, two-tailed t-test with Welch’s correction). This result is consistent with a role for DUSP6 in regulating EGFR-KRAS-ERK signaling^12, 15-19^. DUSP6 RNA was also present at higher levels in LUADs with *EGFR* or *KRAS* mutations than in tumors without such mutations in an independent collection of 83 tumors collected at the BC Cancer Agency (p=0.004), confirming the findings derived from the TCGA dataset (Fig 2B). Furthermore, DUSP6 was more abundant in EGFR/KRAS mutant LUADs than in normal lung tissue (p<0.0001) whereas no significant differences in DUSP6 levels were observed between normal lung tissue and tumors without mutations in either of these two genes (p=0.64) (Fig 2C).

**Figure 2:**
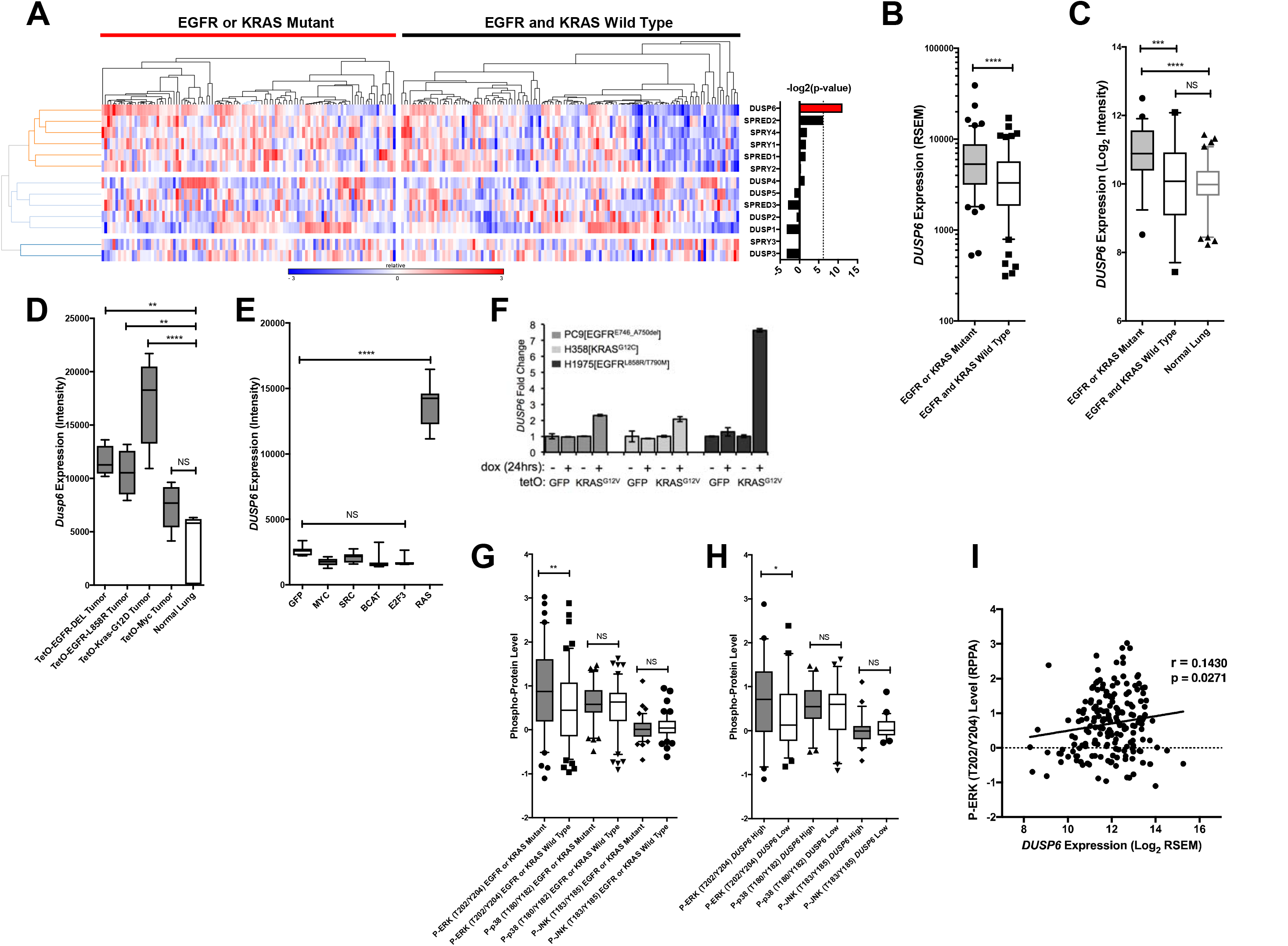
DUSP6 is the only negative feedback regulator significantly up-regulated in LUAD tumors with KRAS or EGFR mutations. (A) Negative feedback regulators differentially expressed between clinical LUADs with or without *EGFR* or *KRAS* mutations. Expression levels for the indicated genes as determined by RNA-seq were compared between LUAD tumors with (n=107, red) and without (n=123, black) *KRAS* and *EGFR* mutations. In the heatmap, red indicates high relative expression and blue, low expression. Significance, as determined by two-tailed unpaired t-test with Bonferroni multiple testing correction, is indicated as the −log_2_(p-value). The significance threshold was set at a p-value <0.01 and is indicated by the dotted line. Only *DUSP6* surpassed this threshold. (B) *DUSP6* is the main negative feedback regulator up-regulated in LUADs with *EGFR* or *KRAS* mutations. Box blots of *DUSP6* expression from samples in A. LUADs with *EGFR* or *KRAS* mutations (n=107) have higher *DUSP6* expression than LUADs with wildtype *KRAS* and *EGFR* (n=123) in the TCGA dataset. (C) Validation of increased *DUSP6* expression in LUADs with mutated *KRAS* or *EGFR*. In an independent internal dataset from the BCCA, LUADs with *EGFR* or *KRAS* mutations (n=54) demonstrated higher expression of *DUSP6* compared to LUADs in which both *EGFR* and *KRAS* were wild-type (n=29) and to normal lung tissues (n=83). (D) *Dusp6* is upregulated in the lungs of mice with tumors induced by mutant *EGFR* or *Kras* transgenes. Tumor-bearing lung tissues from mice expressing *EGFR* or *Kras* oncogenes produce higher levels of Dusp6 RNA than do normal lung controls or tumor-bearing lungs from mice with a *MYC* transgene. (E) Increased DUSP6 RNA is specific to cells with oncogenic signaling through RAS. Human primary epithelial cells expressing a *HRAS* oncogene (n=10 biological replicates) express *DUSP6* at higher levels than control cells producing GFP (n=10 biological replicates) whereas cells expressing known oncogenes other than *RAS* genes do not. (F) DUSP6 RNA levels increase in PC9, H358 and H1975 cells expressing mutant KRAS. Dox was added to induce either *GFP* or the *KRAS*^G12V^ oncogene for 24 hours; DUSP6 RNA was measured by qPCR. (G-I) DUSP6 expression is associated with P-ERK levels. (G) LUADs with *EGFR* or *KRAS* mutations (n=107) have higher P-ERK levels, but not P-p38 or P-JNK levels, than LUADs with wildtype *KRAS* and *EGFR* (n=123) in the TCGA dataset. H) LUADs with the highest *DUSP6* RNA levels (n=46) demonstrated higher P-ERK levels, but not P-p38 or P-JNK levels, than LUADs with the lowest *DUSP6* RNA levels (n=46). I) *DUSP6* RNA levels correlate with the levels of P-ERK in LUADs (n=182). *p<0.05, **p<0.01, ***p<0.001, ****p<0.0001, NS=Not Significant.

To ascertain whether *DUSP6* is up-regulated specifically in tumors driven by mutant KRAS or mutant EGFR signaling rather than in tumors associated with activation of other oncogenic pathways, we measured *DUSP6* in experimental systems driven by the activation of various oncogenes. In transgenic mouse models of lung cancer, *DUSP6* was present at significantly higher levels in the lungs of mice bearing tumors driven by mutant *EGFR* or *KRAS* transgenes than in normal mouse lung epithelium (Fig 2D). In contrast, *DUSP6* levels were not significantly different in lungs from mice with tumors driven by MYC and in normal mouse lung tissue. (Fig 2D). Similarly, increased levels of DUSP6 mRNA were observed in primary human epithelial cells only when they were transduced with mutant *RAS* genes, but not with a variety of other oncogenes or with plasmids encoding GFP (p<0.0001) (Fig 2E). Lastly, our LUAD cell lines engineered to express KRAS^G12V^ in response to dox showed an increase in *DUSP6* that correlated with augmented phosphorylation of ERK and cell toxicity (Fig 2F). Together, these findings suggests that DUSP6 is a critical negative feedback regulator activated in response to oncogenic signaling by mutant RAS or EGFR in LUAD.

In our previous study^5^ (see also Figs. 1 and S1), we found that co-induction of oncogenic KRAS and EGFR activated not only ERK, but also JNK and p38 MAPK pathways at later times. To investigate whether DUSP6 is up-regulated in response specifically to phosphorylation of ERK or also in response to phosphorylation of JNK and p38, we assessed the relationship of amounts of DUSP6 mRNA in tumors with levels of P-ERK, P-JNK and P-p38 proteins as determined for TCGA^14^, using the Reverse Phase Protein Array (RPPA). LUADs with a *KRAS* or an *EGFR* mutation contained significantly higher levels of P-ERK – but not P-JNK or P-p38 – than did tumors without those mutations, consistent with a role for these oncogenes in ERK activation (Fig 2G). Furthermore, tumors with high *DUSP6* RNA have relatively high amounts of P-ERK but not of P-JNK or P-p38 (Fig 2H). Lastly, there is a positive correlation between P-ERK levels and *DUSP6* mRNA in LUAD (Fig 2I), whereas no such association was observed between *DUSP6* mRNA and P-JNK or P-p38 (data not shown). Together, these observations support the proposal that DUSP6 is produced in response to activation of ERK and that it serves as a major negative feedback regulator of ERK signaling in LUAD, buffering the potentially toxic effects of ERK hyperactivation.

### Knockdown of DUSP6 elevates P-ERK and reduces viability of LUAD cells with either KRAS or EGFR oncogenic mutations

If DUSP6 is a negative feedback regulator of RAS signaling through ERK, then inhibiting the enzymatic function of DUSP6 in LUAD cell lines driven by oncogenic KRAS or EGFR should cause hyperphosphorylation and hyperactivity of ERK, producing a signaling intensity that causes cell toxicity, as observed when we co-express mutant KRAS and EGFR. Consistent with this prediction, introduction of *DUSP6*-specific siRNA into PC9 cells knocked down DUSP6 RNA levels and reduced the number of viable cells to levels similar to those observed when *EGFR*, the driver oncogene, was itself knocked down (Fig 3A). siRNAs for both *DUSP6* and *EGFR* decreased DUSP6 RNA levels, further confirming that transcriptional activation of DUSP6 occurs downstream of EGFR signaling. However, while it was anticipated that knockdown of EGFR would diminish the numbers of viable cells by reducing levels of P-ERK, cells in which DUSP6 was knocked down with siRNAs also displayed reduced P-ERK levels at the same time, five days after transfection, contrary to the expected increase in phosphorylation of ERK.

**Figure 3:**
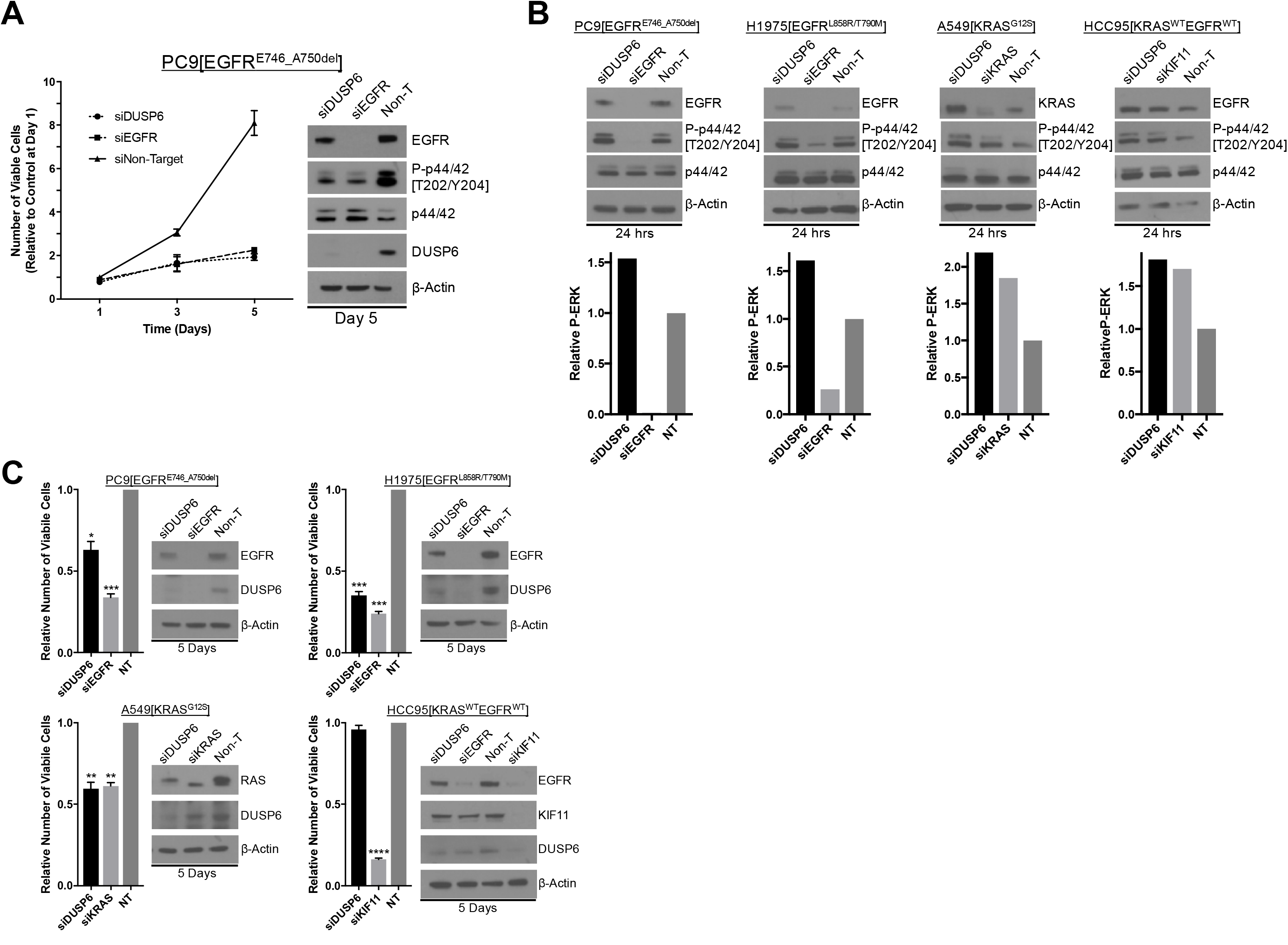
Knockdown of DUSP6 increases P-ERK and selectively inhibits LUAD cell lines with KRAS or EGFR mutations. (A) Interference with *DUSP6* RNA induces toxicity in PC9 cells. Pooled siRNAs for *DUSP6, EGFR* or a non-gene targeting control (Non-T) were transfected into PC9 cells (carrying an *EGFR* mutation) on day 0 and day 3, and the viability of cells in each condition was measured by Alamar blue at the indicated time points and scaled to the Non-T condition at day 1 to measure the relative change in viability. Experiments were done in biological triplicate with the average values presented +/- SEM. Western blots were performed at the endpoint of the assay (day 5) to confirm reduced amounts of DUSP6 protein and measure ERK and P-ERK (p42/44 and P-p42/44, respectively). (B) Interference with *DUSP6* RNA acutely increases P-ERK levels. DUSP6 was knocked down in PC9 and H1975 cells (*EGFR* mutants), A549 cells (*KRAS* mutant), and HCC95 cells (*KRAS* and *EGFR* wild-type); levels of ERK and P-ERK were measured by Western blot 24 hours later. Relative P-ERK levels (ratio of phosphorylated to total levels normalized to actin) were determined by dosimetry and compared to the non-targeting control (NT) to quantify the relative increase after DUSP6 knockdown. (C) Interference with *DUSP6* RNA inhibits LUAD cell lines with activating mutations in genes encoding components of the EGFR/KRAS signaling pathway. Viability of cells 5 days after knockdown of DUSP6 or knockdown of positive controls (EGFR, KRAS or KIF11) was assessed by Alamar blue and compared to the non-targeting controls to determine relative changes. Experiments were done in biological triplicate with the average values presented +/- SEM. Westerns indicating target gene knockdown at Day 5 are also displayed. *p<0.05, **p<0.01, ***p<0.001, ****p<0.0001, NS=Not Significant.

To determine whether an initial, transient increase in P-ERK occurred after DUSP6 knockdown, preceding the observed reduction in viable cells, we measured P-ERK in a panel of cell lines with and without mutations in *EGFR* or *KRAS* 24hrs after addition of DUSP6 siRNA. In all cell lines assessed, there was a small but consistent early increase (~1.5 fold) in P-ERK in cells that received DUSP6 siRNA, compared to non-targeting siRNA controls (Fig 3B). Furthermore, within 5 days, knockdown of DUSP6 reduced the numbers of viable cells in the LUAD lines with activating *KRAS* or *EGFR* mutations (PC9, H1975 and A549), but not in a cell line with no known activating mutations affecting the EGFR-KRAS-ERK pathway (HCC95) (Fig 3C). These data demonstrate that inactivation of DUSP6, a target-specific phosphatase, increases P-ERK and reduces viable cells if they contain an oncogenic *KRAS* or *EGFR* mutation.

### Pharmacological inhibition of DUSP6 reduces the number of viable LUAD cells bearing mutations that activate the ERK pathway

The results presented thus far suggest that LUAD cells with mutations in *KRAS* or *EGFR* depend on DUSP6 to attenuate p-ERK for survival, offering a potentially exploitable vulnerability that could be useful therapeutically. However, efficiently inhibiting DUSP6 with siRNA is difficult, as inactivation of DUSP6 leads to increased p-ERK and a subsequent increase in *DUSP6* mRNA. As *DUSP6* mRNA rises, more siRNA is required to sustain the reduction of DUSP6. Based on this negative feedback cycle, we reasoned that pharmacological inhibition of DUSP6 protein would be more effective. A specific small molecule inhibitor of DUSP6, (E)-2-benzylidene-3-(cyclohexylamino)-2,3-dihydro-1H-inden-1-one (BCI), was identified through an *in vivo* chemical screen for activators of fibroblast growth factor signaling in zebrafish^20, 21^. BCI is an allosteric inhibitor of DUSP6, binding near the active site of the phosphatase, inhibiting its catalytic activation after binding to its substrate, ERK^20^. BCI also selectively inhibits DUSP1, which, like DUSP6, has catalytic activity dependent on substrate binding, but not other DUSPs that are constitutively active. However, as demonstrated in Fig. 2A, *DUSP1* is not significantly up-regulated in LUADs with *EGFR* or *KRAS* mutations, suggesting that DUSP6 could be the main target of BCI in this context.

We tested 11 lung cancer cell lines - 8 with a *KRAS* or *EGFR* mutation and 3 with no known activating mutations in these genes – with a dosing strategy covering the previously determined active range of the drug ^22^. We predicted that cancer lines with mutations in *KRAS* or *EGFR* would be more sensitive to the potential effects of BCI treatment on numbers of viable cells, since DUSP6 would be required to restrain the toxic effects of P-ERK in these cells. Our findings were consistent with this prediction (Fig 4A-B). The cell lines fell into three categories of sensitivity: 1) the most sensitive lines, with IC50s between 1-3uM and with >90% loss of viable cells at 3.2uM, all harbored *KRAS* or *EGFR* mutations; 2) the one line with intermediate sensitivity, H1437 (IC50>4uM), contains an activating mutation in *MEK* (Q56P); and 3) the relatively insensitive lines (IC50s≥5uM) lack known mutations affecting the EGFR-KRAS-ERK signaling pathway. The insensitive cell lines did not demonstrate the marked (>90%) reduction in numbers of viable cells observed with the sensitive cell lines. Together, these data suggest that pharmacological inhibition of DUSP6 specifically kills cells with *EGFR* or *KRAS*-mutations.

**Figure 4:**
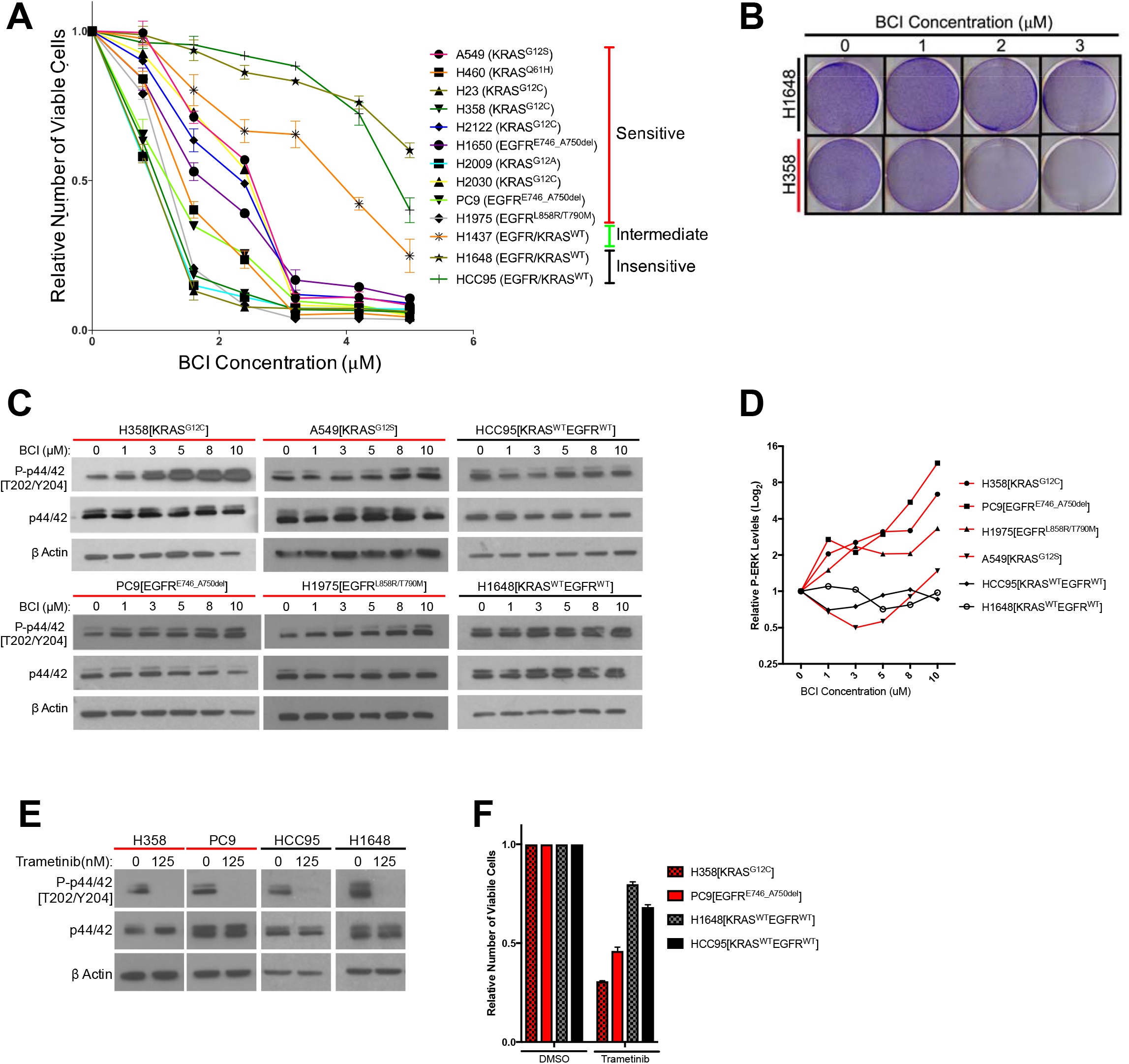
Treatment with the DUSP6 inhibitor BCI selectively kills LUAD cell lines with KRAS or EGFR mutation, implying a dependence on ERK-mediated signaling. (A-B) BCI treatment induces toxicity specifically in lung cancer cell lines with mutations in genes encoding components in the EGFR-KRAS-ERK pathway. (A) Eleven lung cancer cell lines were treated with increasing doses of BCI for 72 hours based on the reported effective activity of the drug^22^. Cell lines could be assigned to three distinct groups: sensitive (red), intermediate (green) and insensitive (black). All sensitive cell lines contained either *EGFR* or *KRAS* mutations; the intermediate and insensitive cell lines were wild-type for genes encoding components of the EGFR-KRAS-ERK signaling pathway (as determined by the Sanger Cell Line Project and the Cancer Cell Line Encyclopedia^42^). Experiments were done in biological duplicate with the average values presented +/- SEM. (B) Crystal Violet stain of cells plated in the indicated doses of BCI or control (0 = 0.1% DMSO) for 72 hours. Sensitive cells with a *KRAS* mutation (H358 cells; denoted with red underlining) show a dramatic decrease in cell number, whereas insensitive cells without oncogenic mutations in genes encoding components of the EGFR-KRAS-ERK pathway (H1648 cells; black underlining) do not. Experiments were done in biological duplicate with a representative image shown. (C) BCI increases P-ERK levels specifically in BCI-sensitive cell lines. Sensitive lines (H358, PC9, H1975 and A549; red underlining) and insensitive lines (HCC95 and H1648; black underlining) were treated with the indicated doses of BCI or vehicle control (0.1% DMSO) for 30 minutes, and the levels of ERK (p44/p42) and P-ERK (P-p44/42 T202/Y204) assessed by Western blot. P-ERK appeared in the sensitive cells at low doses of BCI doses, but P-ERK levels did not increase in the insensitive cells at the tested doses of BCI. (D) Dosimetry plots from the experiment shown in panel (C). (E-F) Cell lines sensitive to BCI are also dependent on P-ERK for survival. BCI-sensitive cells with oncogenic mutations in *EGFR* or *KRAS* (PC9 and H358, respectively; red underlining) and BCI-insensitive cells (H1648 and HCC95; black underlining) were treated with the indicated doses of the MEK inhibitor Trametinib for 72 hours; viable cells were measured with Alamar blue and compared to cells receiving the vehicle control (0 = 0.1% DMSO). (E) Treatment with Trametinib decreased P-ERK levels as determined by western blot. (F) The reduction in PERK corresponded to a greater decrease in viable BCI-sensitive cells (red coloring), compared to BCI-insensitive cell lines (black coloring).

### P-ERK levels increase in LUAD cells after inhibition of DUSP6 by BCI, and P-ERK is required for BCI-mediated toxicity

Based on findings in the preceding section, we predicted that BCI-mediated inhibition of DUSP6 would increase P-ERK to toxic levels, similar to the effects of co-expressing mutant *KRAS* and *EGFR*. To test this proposal, we measured total ERK and P-ERK after BCI treatment in sensitive and insensitive cell lines. A subset of the most sensitive cell lines, H358 (KRAS mutant) and PC9 and H1975 (EGFR mutants), demonstrated a large, dose-dependent increase in P-ERK in response to BCI treatment, with appreciable increases observed even at the lowest doses tested (1uM) (Fig 4C-D). Likewise, another sensitive cell line, A549 (KRAS mutant), demonstrated an increase in P-ERK, albeit at higher BCI concentrations, consistent with a less acute BCI sensitivity (Fig 3C, 4C-D). Conversely, BCI did not induce increases in P-ERK in the insensitive cell lines HCC95 and H1648, even at the highest levels of BCI (10uM) (Fig 4C-D). Importantly, cell lines sensitive to BCI were also dependent on sustained P-ERK signaling for survival, as the MEK inhibitor Trametinib, while effectively reducing P-ERK in all cell lines, reduced cell viability to a greater degree in BCI sensitive lines (H358 and PC9) compared to BCI insensitive lines (H1648 and HCC95; Fig 4E-F). Thus, the oncogenic mutation profile and dependency on activation of the EGFR-RAS-ERK pathway correlates with dependence on DUSP6 activity. These correlations are likely to reflect the central significance of P-ERK as a determinant of cell growth and viability.

To test this relationship further, we predicted that stimulating the EGFR-RAS-ERK pathway in a BCI-insensitive cell line would make the cells more dependent on DUSP6 activity and more sensitive to BCI. Using HCC95 cells, which express relatively high levels of wild-type EGFR (Fig 5B), we provided exogenous EGF (100ng/mL) for one week to stimulate phosphorylation and activation of ERK and then treated the cells with increasing doses of BCI to inhibit DUSP6. After pretreatment with EGF, 3uM BCI reduced the number of viable HCC95 cells by approximately 40% compared to the control culture that did not receive EGF. This outcome implies that EGF makes HCC95 cells dependent on DUSP6 activity, as also observed in cell lines with *EGFR* or *KRAS* mutations (Fig 4A). Indeed, we showed that EGF increased the levels of both P-EGFR and PERK in HCC95 cells, confirming activation of the relevant signaling pathway (Fig 5B and 5C). In addition, BCI further enhanced the levels of P-ERK, especially in the EGF-treated cells, with dose-dependent increases; these findings are similar to those observed in cell lines with *EGFR* or *KRAS* mutations (Fig 4C and 4D). Taken together, these findings suggest that LUAD cells with *KRAS* or *EGFR* mutations are sensitive to BCI because the drug acutely increases P-ERK beyond a tolerable threshold in a manner analogous to the synthetic lethality we previously described in LUAD lines after co-expression of mutant KRAS and EGFR^5^.

**Figure 5:**
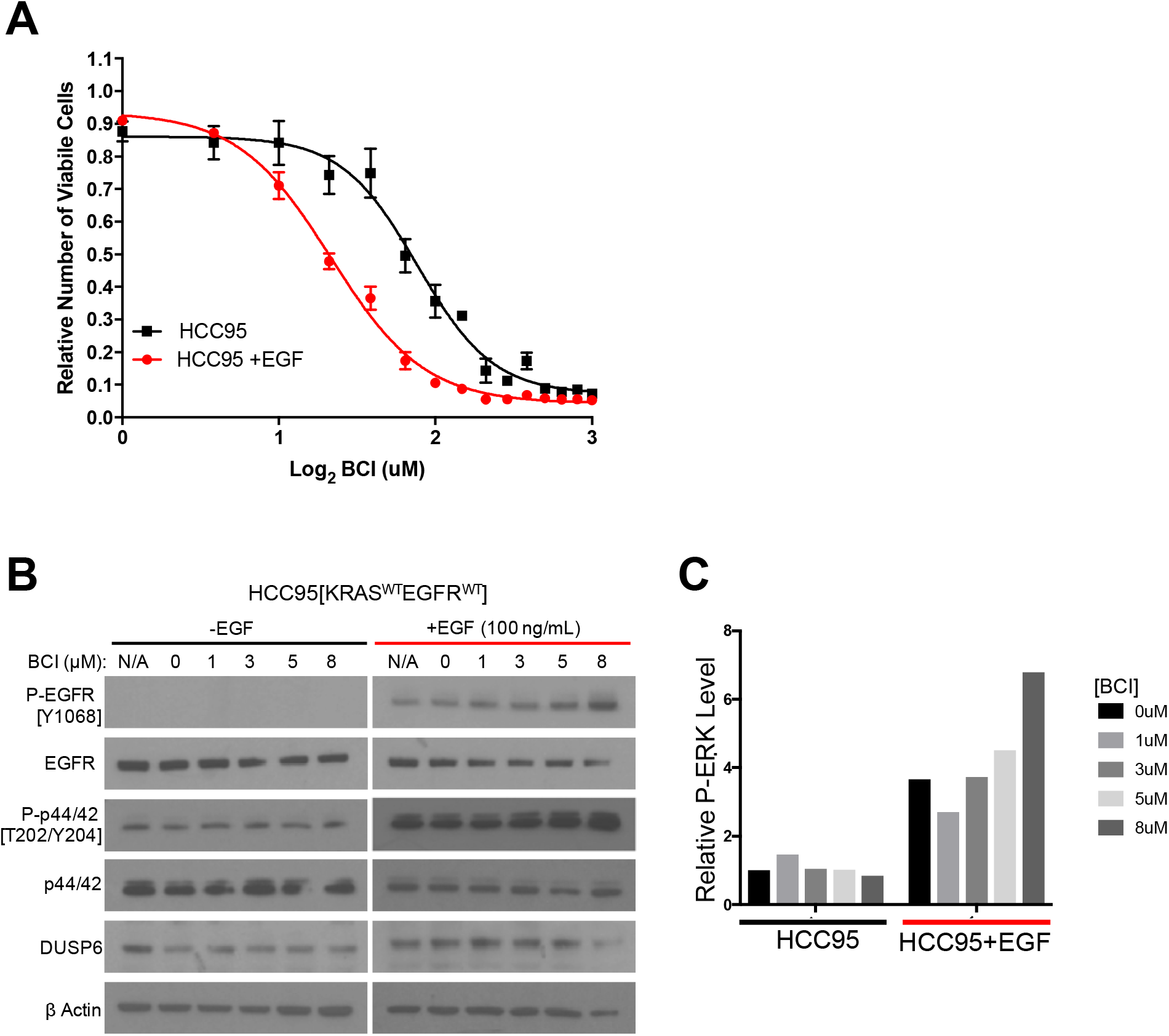
EGF-mediated activation of ERK signaling leads to dependence on DUSP6. (A) Increase of P-ERK promotes sensitivity of lung cancer cell lines without *KRAS* or *EGFR* mutations to BCI treatment. BCI-insensitive HCC95 cells were treated with 100ng of EGF for 7 days and then treated with escalating doses on BCI with continued EGF treatment. Viable cells were measured 72 hours later by Alamar blue and compared to the vehicle controls (in 0.1% DMSO) to assess the relative change in viable cell numbers. Experiments were done in biological triplicate with the average values presented +/- SEM. The EGF-treated cells (red line) showed increased sensitivity (decreased viable cells at lower BCI conditions) than those without EGF treatment (black line). (B-C) EGF increases P-ERK in HCC95 cells. (B) Cells were treated or not treated with EGF and increasing doses of BCI; levels of the indicated proteins were assessed in cell lysates by Western blotting. EGF increased the levels of P-EGFR and P-ERK, which were further elevated by BCI. (C) Relative P-ERK levels (ratio of phosphorylated to total levels normalized to actin) were determined by dosimetry and compared to the vehicle controls (0 BCI = 0.1% DMSO) to quantify the relative increase after BCI treatment from the gels in B.

## Discussion

The pattern of mutual exclusivity observed with mutant *EGFR* and mutant *KRAS* genes in LUAD is a consequence of synthetic lethality, not pathway redundancy; co-expression of these oncogenes is toxic to cells, resulting in loss of viable cells^5, 8^. But why are cells unable to tolerate the combination of these two oncogenes? And what are the biochemical mechanisms by which the toxicity is controlled?

Our efforts to answer these questions have led us to the conclusions that the toxicity is mediated through the hyperactivity of phosphorylated ERK1/2 and that inhibition of DUSP6 can re-create the toxicity through the role of this phosphatase as a negative regulator of ERK1/2. Several results reported here support these conclusions: (i) the previously reported toxicity that results from co-expression of mutant *EGFR* and mutant *KRAS* is accompanied by an early increase in the phosphorylation of ERK1/2, and the effects can be attenuated by inhibiting MEK (which phosphorylates and activates ERK1/2) or by reducing ERK levels with inhibitory RNAs; (ii) DUSP6, a phosphatase known to be a feedback inhibitor of ERK activity, is present at relatively high levels in LUADs with *EGFR* and *KRAS* mutations; and (iii) inhibition of DUSP6, either by introduction of siRNAs or by treatment with the drug BCI, reduces the number of viable LUAD cells with *EGFR* or *KRAS* mutations or of BCI-resistant cells exposed to EGF.

Taken in concert, these findings support a general hypothesis about cell signaling. Activation of a biochemical signal from a critical node, such as ERK, in a signaling pathway must rise to a certain level to drive neoplastic changes in cell behavior; if signal intensity falls below that level, the cells may revert to a normal phenotype or initiate cell death as a manifestation of what is often called “oncogene addiction”^23-27^. Conversely, if the intensity of signaling rises to exceed a higher threshold, the cells may display a variety of toxic effects, including senescence, vacuolization, or apoptosis^5, 28-32^. In this model, two approaches to cancer therapy can be envisioned: (i) blocks to signaling that reverse the oncogenic phenotype or induce the apoptosis associated with oncogene addiction, or (ii) enhancements of signaling that cause selective toxicity in cells with pre-existing oncogenic mutations, a form of synthetic lethality that depends on changes that produce a gain rather than a loss of function. The former is exemplified by using inhibitors of EGFR kinase activity to induce remissions in LUAD with EGFR mutations^33-35^. Based on the findings presented here, the latter strategy might be pursued by using inhibitors of DUSP6 to block its usual attenuation of signals emanating from activated ERK1/2.

Several factors are likely to determine the threshold for producing the cell toxicity driven by hyperactive signaling nodes, such as ERKs, in cancer cells. These factors are likely to include allele-specific attributes of oncogenic mutations in genes such as *KRAS*^36^ and *BRAF*^36-38^; the cell lineage in which the cancer has arisen^22, 37, 39^; the levels of expression of mutant cancer genes^32, 38, 40, 41^; the co-existence of certain additional mutations^42^; and the multiple proteins that negatively regulate oncogenic proteins through feedback loops, such as MIG6 on EGFR^41, 43, 44^, GAPs on RAS proteins^45, 46^, or SPROUTYs and DUSPs on kinases downstream of RAS^18, 22, 39^. All such factors would need to be considered in the design of therapeutic strategies to produce signal intensities that are intolerable specifically in cancer cells.

The findings presented here, as well as recent results from others^22^, support several underlying features of a therapeutic strategy based on inordinate signaling activity involving RAS proteins: that the activity of ERK needs to be actively controlled in cancer cells of diverse tissue origins; that hyperactivation of ERK can be deleterious to cells; and that inhibition of negative regulators like DUSP6 can create a toxic cellular state. This leads to the hypothesis that cancer cells *dependent* on ERK signaling have an active RTK-RAS-RAF-MEK pathway that produces levels of activated (phosphorylated) ERK1/2 that *require* attenuation. In other words, ERK-dependent tumor cells, including cancers driven by mutant RTK, RAS, BRAF, or MEK proteins, will have a vulnerability to hyperactivated ERK and that vulnerability can potentially be exploited by inhibition of feedback regulators like DUSP6.

This hypothesis is particularly attractive in view of the frequency of *RAS* gene mutations in human cancers and the difficulties of targeting mutant RAS proteins^47-49^. Because DUSP6 directly controls the activities of ERK1 and ERK2, rather than proteins further upstream in the signaling pathway, it appears to be well-situated for controlling both the signal delivered to ERK through the activation of RAS and the signal emitted by phosphorylated ERK. Inhibition of other DUSPs, like DUSP5, that regulate ERK1 and ERK2 may create similar vulnerabilities and should be explored^18, 50^. These ideas should provoke searches for inhibitors of DUSPs and other feedback inhibitors of this signaling pathway, as well as experiments that better define the downstream mediators and the consequences of non-attenuated ERK signaling.

## Materials and Methods

### Cell lines and culture conditions

PC9 (PC-9), H358 (NCI-H358), H1975 (NCI-H1975), H1648 (NCI-H1648), A549 and HCC95 cells were obtained from either American Type Tissue Culture (ATCC) or were a kind gift from Dr. Adi Gazdar. For experiments involving doxycycline inducible constructs, cells were maintained in RPMI-1640 medium (Lonza) supplemented with 10% Tetracycline-free FBS (Clontech) or FBS that was tested to be Tet-free (VWR Life Science Seradigm), 10mM HEPES (Gibco) and 1mM Sodium pyruvate (Gibco). For other experiments, cells were grown in RPMI-1640 medium (Thermo Fisher) supplemented with 10% FBS (Sigma), 1% Glutamax (Thermo Fisher) and Pen/Strep (Thermo Fisher). Cells were cultured at 37°; air; 95%; CO2, 5%. Where indicated, doxycycline hyclate (Sigma-Aldrich) was added at the time of cell seeding at 100 ng/ml. Trametinib (Selleckchem), SCH772984 (Selleckchem), Dual Specificity protein phosphatase 1/6 inhibitor (BCI) (Calbiochem), and EGF recombinant human protein solution (Thermo Fisher) were added at the time of cell seeding at the indicated doses.

### Plasmids and generation of stable cell lines

Plasmids used were identical to those described in a prior publication^5^. In brief, DNAs encoding mutant KRAS or GFP were cloned into pInducer20, a vector that carries a tetracycline response element for dox-dependent gene control and encodes rtTA, driven from the UbC promoter^9^. Lentivirus was generated using 293T cells (ATCC), psPAX2 #12260 (Addgene, Cambridge, MA) and pMD2.G (Addgene plasmid#12259). Polyclonal cell lines (H358-tetO-GFP, H358-tetO-KRAS^G12V^, PC9-tetO-GFP, H1975-tetO-GFP) and single cell-derived clonal cell lines (PC9-tetO-KRAS^G12V^, H1975-tetO-KRAS^G12V^) were used. pLKO.1-based lentiviral vectors were used to establish cells stably expressing shRNAs for the indicated genes. Knockdown was achieved using two independent shRNAs targeting *ERK1* (noted in text as A4 and A5) or *ERK2* (noted in text as G6 and G7) RNAs. An shRNA targeting GFP was used as a control. shRNA constructs were kindly provided by J. Blenis, Weill Cornell Medicine. Lentivirus was generated using 293T cells as above. After transduction, polyclonal cells were selected with puromycin and maintained as a stable cell line.

### Measurements of protein levels

Cells were lysed in RIPA buffer (Boston Bioproducts) containing Halt protease and phosphatase inhibitor cocktail (Thermo Fisher). For experiments involving dox-inducible constructs, lysates were cleared by centrifugation, and protein concentration determined by Pierce BCA protein assay kit (Thermo Fisher). Samples were denatured by boiling in loading buffer (Cell Signaling). 20 μg of lysates were loaded on 10% MiniProtean TGX gels (Bio-Rad), transferred to Immun-Blot PVDF membranes (Bio-Rad), blocked in TBST (0.1% Tween-20) and 5% milk. For all other experiments, samples were denatured by boiling in loading buffer (BioRad) and 25 μg of lysates were loaded on 4-12% Bis-Tris gradient gels (Thermo Fisher), run using MOPS buffer, transferred to Immobilon-P PVDF membranes (Millipore) and blocked in TBST (0.1% Tween-20)/5% BSA (Sigma).

Primary incubation with antibodies was performed overnight at 4° in 5% BSA, followed by appropriate HRP-conjugated secondary antisera (Santa Cruz Biotechnology) and detected using ECL (Thermo Fisher).

Antibodies were obtained from Cell Signaling and raised against the following proteins: phospho p-38 (4511), p38 (8690), p-p44/p42 (ERK1/2) (9101), p44/p42 (ERK1/2) (4695), p-SAPK/JNK (4668), SAPK/JNK (9252), P-EGFR (3777, 2234), EGFR (2232), KRAS (8955), a-Tubulin (3873) and b-Actin (3700, 4970). Additionally, we used an antibody against GFP (A-21311, Thermo Fisher).

For 24 hour time course experiments, 100,000 cells (PC9, H1975) or 500,000 cells (H358) per well were seeded in a 6-well plate and stimulated with dox or dox and drug. For 5-day experiments, 25,000 cells were seeded in 6-well format.

For proteome profiler array, 200 ug of total lysate was incubated on membranes in the A/B set (ARY003B, R&D Systems) and processed according to protocol (R&D Systems). Film exposures were scanned and spot density quantified using Image Studio Lite (Licor). Data were plotted in Microsoft Excel.

For western blots with BCI and Trametinib, cells were seeded to achieve 80% confluency 18 hours post seeding. Medium was aspirated and replaced with antibiotic-free medium containing drug at indicated concentrations and incubated for 30 minutes. Cells were lysed and protein levels assessed as stated above. Quantification of western blot images was performed using ImageJ software. Scanned files were saved in TIFF format, and background was subtracted from all images. Rectangle tool was used to fully encompass each separate band. Rectangles and bands were assigned lanes and histogram plots were generated based on each lane. Each histogram was enclosed using a straight line across the bottom and the “magic wand” tool generated a value for area of histogram. These values were exported to and assessed using Excel and Graphpad Prism software.

### Measurements of viable cells

For experiments with dox-inducible constructs, cells were seeded into media containing doxycycline (100 ng/ml) and/or drug (Trametinib, SCH772984). Media (with or without doxycycline or drug) were replenished every 3 days during the 7 days. At indicated time points, medium was aspirated and replaced with medium containing Alamar Blue (Thermo Fisher). Fluorescence intensities from each well were read in duplicate on a FLUOstar Omega instrument (BMG Labtech), and data plotted in Microsoft Excel. Cells were seeded in triplicate in 24-well format at 1,000 cells/well (PC9 or H1975 derivatives) or 5,000 cells/well (H358 derivatives). For other experiments, cells were grown in 6-well plates, Alamar Blue added, and intensities measured for each well in quadruplicate using a Cytation 3 Multi Modal Reader with Gen5 software (BioTek).

For crystal violet assays, cells were seeded to achieve 80-90% confluency of 80 – 90% at the end point in the absence of drug treatment. 18 hours later, medium was aspirated and replaced with medium containing drug. Cells were incubated for 72 hours, washed with PBS and Crystal Violet solution (Sigma) was added and incubated for 2 minutes before washing again with PBS and imaging.

### Genomic datasets and analyses

RNA-Seq (RSEM) data for EGFR-KRAS-ERK pathway phosphatases (DUSP1-6, SPRED1-3, SPRY1-4) along with corresponding mutational data for *EGFR* and *KRAS* for 230 lung adenocarcinoma tumors from The Cancer Genome Atlas^14^ were downloaded from cBioPortal (<u>http://www.cbioportal.org/</u>)^51, 52^. Expression of each gene was compared between tumors with *KRAS* or *EGFR* mutations and those without, using an unpaired T-Test. Resulting p-values were adjusted for multiple comparisons using a Bonferroni correction and the −Log_2_ value plotted as an indication of significance. Expression values for each gene were also median normalized across the samples and plotted using GENE-E software (Broad Institute) as a heat map. Reverse phase protein array (RPPA) data (replicate-base normalized^53^) for 182/230 tumors were downloaded from the UCSC Cancer Genomics Browser. Levels of MAPKPT202Y204, P38PT180Y18 and JNKPT183Y185 were compared between samples with a *KRAS* or *EGFR* mutation and those without, using the unpaired T-test. Likewise, samples were separated into groups with high and low DUSP6 expression levels, based on the highest and lowest *DUSP6* expression quartiles; MAPKPT202Y204, P38PT180Y18 and JNKPT183Y185 levels were compared between the groups as above. Lastly, MAPKPT202Y204 levels from RPPA (RBN values) were correlated with *DUSP6* expression (Log_2_ RSEM values), and the Pearson correlation coefficient and p-value determined.

*DUSP6* RNA levels were also compared between tumors with and without *EGFR* or *KRAS* mutation in 83 tumors and matched normal lung tissues from the BC Cancer Agency (BCCA) and deposited in the Gene Expression Omnibus (GSE75037) as described above. Similarly, *DUSP6* expression was compared between human epithelial cells expressing various oncogenes or GFP control (GSE3151). Lastly, Affymetrix Mouse Genome 430 2.0 Arrays were used to profile the lung from genetically engineered mouse models of lung cancer with and without the expression of different driver oncogenes (*CCSP-rtTA;TetO-EGFR^ΔL747-S752^, CCSP-rtTA;TetO-EGFR^L858R^, CCSP-rtTA;TetO-Kms^G12D^* and *CCSP-rtTA;TetO-MYC*)^54-56^ and levels of DUSP6 compared using the T-Test as described above.

### siRNA transfections

For the time course experiments, 50,000 cells (PC9) per well were seeded in a 6-well plate. For the endpoint experiments, 50,000 cells (PC9) or 75,000 cells (1975, A549, HCC95) per well were seeded. Cells were then transfected with ON-TARGETplus siRNA pools (Dharmacon) against the following targets as previously described^57^--- EGFR (L-003114-00-0010), KIF11 (L-003317-00-0010), KRAS (L-005069-00-0010), DUSP6 (L-003964-00-0010)—as well as a non-targeting control (D-001810-10-20). For consistent transfection efficiency across experiments, 10uL of 20uM siRNA pool was added in 190uL of OptiMEM (Life Technologies) and 5uL of Dharmafect was added in 195uL of OptiMEM (Life Technologies) at room temperature. The siRNA and Dharmafect suspensions were mixed and incubated for 20 minutes prior to transfection. Media was changed 24 hours after transfection. For sustained knockdown of targets, transfections were conducted on Day 0 and again on Day 3. Viable cells were measured using Alamar Blue as described above. For the time course experiment, cell viability was determined on Day 1, Day 3 (prior to second transfection) and Day 5 or only on Day 5. Results were compared between each siRNA and non-targeting control using a one-sample t-test as previously described^57^.

### BCI dose-response treatments

Dose-response curves for BCI were established using a modified version of the protocol previously described^57^. Briefly, cells were seeded in quadruplicate at optimal densities into 96-well plates containing media with and without BCI at indicated doses in 0.1% DMSO. Viable cells were measured 72-hours later with Alamar Blue as described above. All experiments were performed in at least biological duplicate and plotted +/- SEM. For HCC95 sensitization assays, cells were cultured with or without 100ng/mL of EGF Recombinant Human Protein Solution (Life Technologies) for 10 days prior to seeding in 96-well plates for BCI dose response assays with or without EGF. The cells were allowed to adhere for 24 hours before treatment with 17 different concentrations of BCI, ranging from 0 to 8uM, with 0.5uM increment doses at 0.1% DMSO concentration. Additionally, 100uM of Etoposide (0.1% DMSO) was added as a positive control for cell death. Cell viability was determined after 72 hours of drug exposure using Alamar Blue. Graphpad Prism software was used to create dose response curves.

### Quantitative RT-PCR

Cells were homogenized and RNA extracted using the RNeasy Mini kit (Qiagen) according to the manufacturer’s instructions. cDNA was prepared using the High-Capacity cDNA Reverse Transcription kit (Thermo Fisher). RT–PCR reactions were carried out using the TaqMan Gene Expression Master Mix (Thermo Fisher) and TaqMan Gene Expression Assays (Thermo Fischer) for *DUSP6* (Hs00169257_m1) and *GAPDH* (Hs99999905_m1). Reactions were run on a QuantStudio6 Real Time PCR system (Thermo Fisher). The ΔΔCt method was used for relative expression quantification using the average cycle thresholds.

## Supplemental Information

**Supplemental Figure 1:**
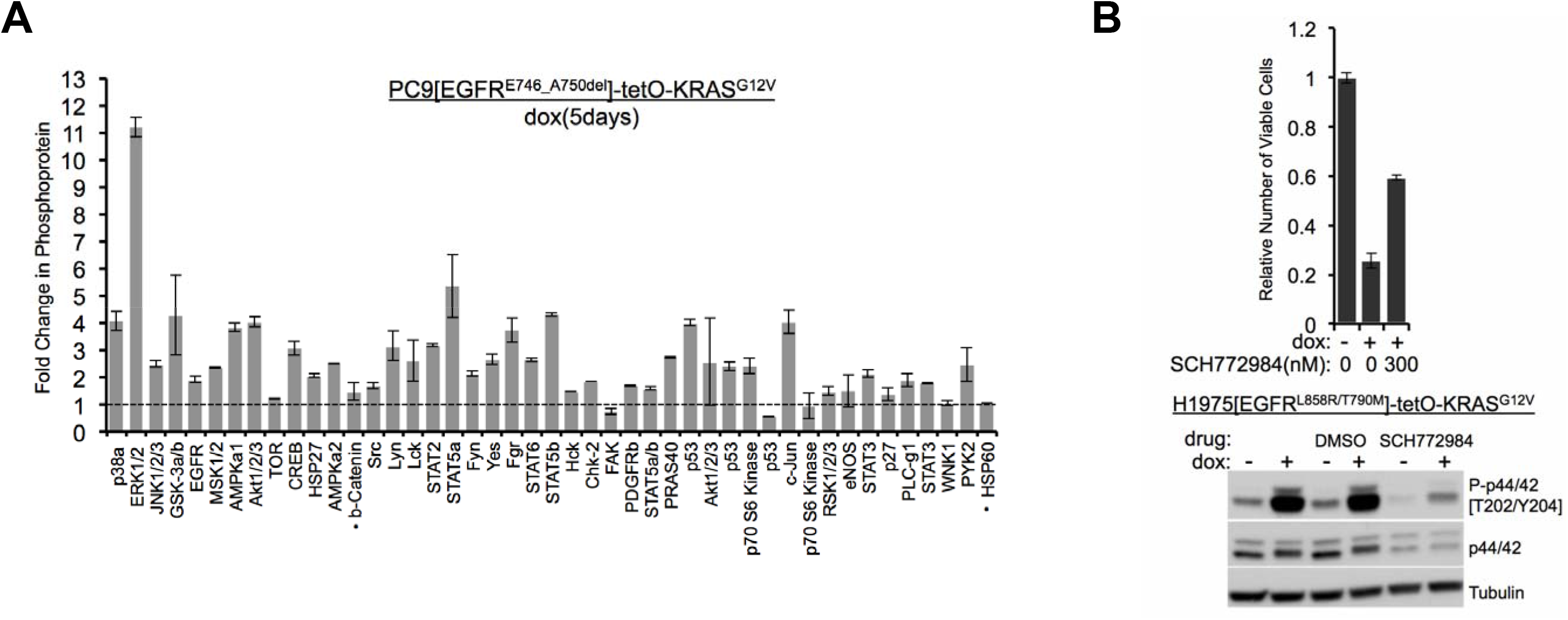
(A). Multiple proteins are phosphorylated after prolonged induction of mutant *KRAS*. Lysates from PC9-tetO-KRAS cells treated or not treated with dox for 5 days were incubated on an array to detect changes in phosphorylation of 43 proteins. HSP60 and b-catenin signals represent total protein content, not phosphoprotein. Fold-changes in dox-treated cells (compared with lysates from untreated cells [set to 1.0, dotted line] are shown from a single antibody array, with error bars derived from duplicate spots on the array. (B) Inhibition of both ERK1 and ERK2 can rescue viability defects in H1975-tetO-KRAS cells. Cells were treated with SCH772984, an ERK inhibitor, and dox for 7 days. The number of viable cells was measured with Alamar blue. Error bars represent standard deviations determined from three samples grown under each set of conditions. Values were normalized to measurements of cells that did not receive either dox or SCH772984. H1975-tetO-KRAS cells were also treated with dox and SCH772984 for 24 hours at the dose conferring a rescue of viable cell numbers (300nM). Lysates were probed for indicated proteins to confirm inhibition of ERK1/2.

## Acknowledgement

We would like to thank Katerina Politi (Yale University) for providing gene expression data from her transgenic mice. We would like to thank members of the Varmus lab for useful discussions and Oksana Mashadova, in particular, for experimental help.

